# Lactate dehydrogenase is associated with cholesterol/lipid metabolism, and fluvastatin plus dipyridamole suppresses canine hemangiosarcoma growth in patient-derived xenograft models

**DOI:** 10.64898/2026.03.03.709271

**Authors:** Tamami Suzuki, Suzune Tanaka, Keika Kishimoto, Takuma Goto, Jumpei Yamazaki, Takashi Kimura, Keisuke Aoshima

**Affiliations:** Laboratory of Comparative Pathology, Department of Clinical Sciences, Faculty of Veterinary Medicine, Hokkaido University, Sapporo, Hokkaido 060-0818, Japan; Translational Research Unit, Veterinary Teaching Hospital, Faculty of Veterinary Medicine, Hokkaido University, Sapporo, Hokkaido 060-0818, Japan; Cancer Research Unit, One Health Research Center, Hokkaido University, Sapporo, Hokkaido 060-0818, Japan

**Keywords:** Drug Therapy, Combination, Gene Expression Profiling, Histones, Neoplasm Transplantation, Neoplasms, Vascular Tissue

## Abstract

Tumor cells commonly exhibit aerobic glycolysis and produce lactate despite oxygen availability. Lactate dehydrogenase (LDH) catalyzes pyruvate-lactate interconversion and regulates intracellular lactate levels. Endothelial cells also depend on glycolysis for ATP production, which prompted us to investigate LDH in canine hemangiosarcoma (HSA), a malignant endothelial tumor. We inhibited LDH with (R)-GNE-140 or sodium oxamate in two canine HSA cell lines (HU-HSA-2 and HU-HSA-3) and generated HU-HSA-3 clones with knockout of *LDHA* or *LDHB* to evaluate the effects of LDH perturbation. (R)-GNE-140 and sodium oxamate suppressed proliferation and reduced global histone lactylation levels in both cell lines. mRNA-sequencing (mRNA-seq) of (R)-GNE-140-treated HU-HSA-2 cells identified cholesterol/lipid metabolism-related gene sets among the top negatively enriched pathways. Representative cholesterol/lipid metabolism genes responded differently depending on cell lines and inhibitors. (R)-GNE-140 decreased these genes in HU-HSA-2 but not HU-HSA-3, whereas sodium oxamate decreased them in HU-HSA-3 with limited effects in HU-HSA-2. In HU-HSA-3, *LDHA* and *LDHB* knockout clones decreased SREBP2 expression and reduced the number of lipid droplets. Fluvastatin, a cholesterol metabolism inhibitor, inhibited HSA cell growth *in vitro* but did not significantly suppress tumor growth in two HSA patient-derived xenograft (PDX) models. In contrast, combined fluvastatin and dipyridamole treatment inhibited proliferation *in vitro* and tumor growth in PDX models. Collectively, these results suggest a context-dependent association between LDH and cholesterol/lipid metabolism in canine HSA cell lines and provide a rationale for further evaluation of combined cholesterol pathway inhibition.

## Introduction

Tumor cells exhibit altered metabolism, including aerobic glycolysis (the Warburg effect), which generates abundant lactate. High lactate concentrations in the tumor microenvironment are associated with aggressive behavior and poor prognosis, and lactate can promote tumor progression through effects on tumor and stromal tissues.^1^ Lactate levels are regulated by lactate dehydrogenase (LDH), which catalyzes the interconversion of pyruvate and lactate while regenerating NAD□ from NADH.^2^ LDH functions as a tetramer composed of LDHA and LDHB subunits, and isoform composition and subcellular localization can vary across tissues and tumors. Notably, nuclear localization of LDHA has been reported in some contexts, supporting potential noncanonical roles linking metabolism to chromatin regulation.^3^ Indeed, lactate is also associated with epigenetic regulation. Histone lactylation is a lactate-derived post-translational modification whose global level correlates with intracellular lactate concentration and can influence transcriptional programs relevant to cancer biology.^4–6^ Beyond glycolysis, tumor cells extensively reprogram lipid metabolism. Cholesterol uptake and biosynthesis support membrane biogenesis and oncogenic signaling,^7^ and tumor cells store excess cholesterol as cholesteryl esters within lipid droplets.^8^ Nutrient-sensing pathways such as the mechanistic target of rapamycin complex 1 (mTORC1) can promote the activation of sterol regulatory element-binding proteins (SREBPs) and lipogenesis.^9^ Statins inhibit HMG□CoA reductase, but their efficacy is often limited by sterol-regulated feedback mediated by SREBP2, a master transcription factor for cholesterol biosynthesis genes.^10,11^ Dipyridamole, an antiplatelet agent and vasodilator, inhibits SREBP2 maturation and can enhance the anti-tumor effect of statins in human cancers.^10–12^

Canine hemangiosarcoma (HSA) is a highly malignant vascular endothelial tumor with poor prognosis. Standard treatment consists of surgical resection followed by adjuvant chemotherapy (most commonly doxorubicin-based), yet the reported 2-year survival rate remains as low as 3.8%.^13,14^ Endothelial cells rely primarily on glycolysis even under aerobic conditions,^15^ suggesting that lactate metabolism may be associated with HSA biology. However, the role of LDH in HSA remains largely unexplored.

In this study, we investigated how LDH contributes to HSA pathogenesis. Our analyses revealed that LDH perturbation profoundly alters cholesterol/lipid metabolism. Therefore, we evaluated pharmacologic inhibition of the cholesterol pathway using fluvastatin with or without dipyridamole in canine HSA cell lines and patient-derived xenograft (PDX) models.

## Materials and Methods

Full materials and methods are found in the supplementary information.

### Cell culture

Two canine HSA cell lines (HU-HSA-2 and HU-HSA-3), isolated from splenic hemangiosarcoma canine patients,^6^ and 293T cells, obtained from RIKEN Bioresource Center, were cultured in Dulbecco’s Modified Eagle Medium (DMEM; #044-29765, FUJIFILM Wako Pure Chemical Corporation, Osaka, Japan) supplemented with 10% fetal bovine serum (FBS; #10270-106, Thermo Fisher Scientific, Waltham, MA, USA) and penicillin-streptomycin (P/S; #168-23191, FUJIFILM Wako) at 37°C with 5% CO_2_.

### Cell line authentication statement

No additional authentication was performed in this study. HU-HSA-2 and HU-HSA-3 cells were established and authenticated in a previous study.^6^ 293T cells were authenticated in RIKEN bioresource center (RCB2202).

### LDHA and LDHB knockout cell generation

293T cells were transfected with 4lentiCRISPRv2 containing designed single guide RNAs (sgRNAs), HIV-gp, and VSV-G using Lipofectamine 3000 with P3000 reagent (#L300015 Thermo Fisher Scientific) in Opti-MEM (#31-985-070, Thermo Fisher Scientific) according to the manufacturer’s instructions. Forty-eight hours after transfection, medium was filtered and collected as viral solutions. Selection was done with medium containing puromycin. Cells were seeded into 96-well plates at a density of 0.5 cells per well to obtain clones.

### Cell proliferation assay

The number of live cells was counted using Countess □ (AMQAX1000, Thermo) or using Cell Counting Kit-8 (CCK-8; #343-07623, Dojindo Laboratories, Kumamoto, Japan) according to the manufacturer’s instructions. Relative viability was calculated by setting control samples as 100%. KyPlot 6.0 software (KyensLab Inc., Tokyo, Japan) was used to generate survival curves.^16^

### Protein extraction and western blotting

SDS lysis buffer was added to cultured cells after washing with phosphate-buffered saline (PBS) twice. Protein concentrations were measured with TaKaRa BCA Protein Assay Kit (#T9300A, Takara Bio Inc., Kusatsu, Shiga, Japan). Proteins were separated on 8%-18% SDS-polyacrylamide gels. Protein-transferred membranes were blocked with 3% skim milk in Tris-buffered saline with 0.05% Tween20 (TBST) for 1 hour at room temperature (RT) and incubated with primary antibodies overnight at 4°C. Membranes were then incubated with the corresponding secondary antibodies. Antibodies used in this study are listed in Table S1.

### mRNA-seq

HU-HSA-2 cells were treated with DMSO or 10 µM (R)-GNE-140 for 72 hours in triplicate. Sequence reads were mapped to ROS_Cfam_1.0 (CanFam 4) using STAR,^17^ and expression levels were estimated using RSEM.^18^ Differential expression analysis was performed using edgeR, and gene expression profiles were analyzed by gene set enrichment analysis (GSEA) v4.1.0.^19,20^ Gene Ontology analysis was conducted using Metascape.^21^

### RNA extraction and reverse transcription quantitative polymerase chain reaction (RT-qPCR)

Total RNA was extracted by using Tripure Isolation Reagent (#11667157001, Roche, Basel, Switzerland), and reverse transcription was performed using PrimeScript Reverse Transcriptase □ (#2690A, Takara Bio) according to the manufacturer’s instructions. qPCR was performed using the cDNAs and primers listed in Table S2. Results were normalized based on the geometric mean of reference genes, which were selected from seven potential internal controls (*RPL32, RPL13A, TBP, YWHAZ, HMBS, B2M,* and *ACTB*) using geNorm software.^22^ Selected reference genes for each experiment are listed in Table S3.

### Lipid droplet staining

For visualization of lipid droplets, scramble, *LDHA*, and *LDHB* knockout HU-HSA-3 clones were fixed with 4% paraformaldehyde for 7.5 minutes at RT, and stained with Lipi-Green (#LD02, Dojindo) diluted in PBS according to the manufacturer’s instructions. Images were acquired using a confocal laser microscope (LSM 800, Carl Zeiss, Oberkochen, Germany). For quantification of lipid droplets, 30 confocal images (one cell per image) were analyzed for each clone and quantified using ImageJ.^23^ Individual cells were manually outlined as regions of interest. Images were converted to 8-bit and thresholded using Li’s method (dark background).^24^ A binary mask was generated, droplets were separated using the watershed algorithm, and particles were quantified with Analyze Particles (size: 0.0005-0.10; circularity: 0-1.00).

### Animal studies

All mouse experiments were performed under the guidelines (protocol number: 20-0083, 21-0062, 25-0062). Female 4- to 8- week-old KSN/Slc mice (*Mus musculus*, Japan SLC, Inc.) were used for experiments. PDX tumor fragments were transplanted subcutaneously in the right flank of mice. Tumor volumes were calculated using the formula: volume = (length × width^2^)/2. When the tumor volume reached 100 mm^3^, treatments were started. For HU-HSAPDX-3 and HU-HSAPDX-5, mice were intraperitoneally injected with fluvastatin (30 mg/kg) every other day. PBS was used as the vehicle control. The treatment duration was up to 30 days or until the tumor volume reached 1 cm^3^. For HU-HSAPDX-6 and HU-HSAPDX-7, mice were orally administered with fluvastatin (50 mg/kg) and/or injected intraperitoneally with dipyridamole (120 mg/kg) daily. Dipyridamole was dissolved in a solvent of 50% (v/v) polyethylene glycol 600 and 2 mg/mL tartaric acid to achieve a final concentration of 5 mg/mL, and this solvent was also used as the vehicle control for dipyridamole. PBS was used as the vehicle control for fluvastatin. The treatment duration was 2 weeks. Mice were euthanized with CO_2_ at the end of treatments.

### Statistical analysis

Statistical analyses were performed with Microsoft Excel (version 2019) and R software (version 4.2.0). For *in vivo* experiments, tumor growth was analyzed with two-way ANOVA. Cell proliferation was analyzed with two-way ANOVA and Dunnett’s test. Statistical comparisons for lipid droplet quantification and qPCR among the three groups (scramble, LDHA knockout, and LDHB knockout clones) were performed using the Kruskal-Wallis test. Dunn’s multiple comparison test with Benjamini-Hochberg correction was subsequently used for post-hoc analysis.

## Results

### LDH is associated with cholesterol/lipid metabolic pathways in HU-HSA-2 and HU-HSA-3 cell **lines**

To investigate the role of LDH activity in HSA cells, we evaluated the effects of (R)-GNE-140, a dual inhibitor of LDHA and LDHB, on cell proliferation and global histone lactylation levels in two HSA cell lines (HU-HSA-2 and HU-HSA-3). Given that lactate can serve as a substrate for histone lactylation, we monitored global histone lactylation levels as a surrogate readout of the downstream effect of LDH inhibition. (R)-GNE-140 significantly suppressed proliferation of both HU-HSA-2 and HU-HSA-3 *in vitro* (Fig. 1A). In HU-HSA-2, LDHA and LDHB protein levels were modestly increased after (R)-GNE-140 treatment, possibly reflecting compensatory upregulation, whereas no obvious change was observed in HU-HSA-3 (Fig. 1B). Global histone lactylation levels were reduced in both cell lines (Fig. 1B). Next, to identify gene expression changes associated with LDH inhibition, we performed mRNA-seq on HU-HSA-2 cells treated with (R)-GNE-140 because HU-HSA-2 showed a more pronounced antiproliferative response. GSEA revealed that multiple cholesterol/lipid metabolism gene sets were among the top 10 negatively enriched pathways (Fig. 1C, D), and Gene Ontology enrichment analysis using Metascape of significantly downregulated genes (≥2-fold; *q* < 0.05) similarly highlighted terms related to cholesterol/lipid metabolism (Fig. 1E). Furthermore, downregulation of representative cholesterol/lipid metabolism genes was validated by RT–qPCR in HU-HSA-2 (Fig. 1F). In contrast, (R)-GNE-140 did not significantly reduce the expression levels of representative cholesterol/lipid metabolism genes in HU-HSA-3 (Fig. 1F).

**Fig. 1.**
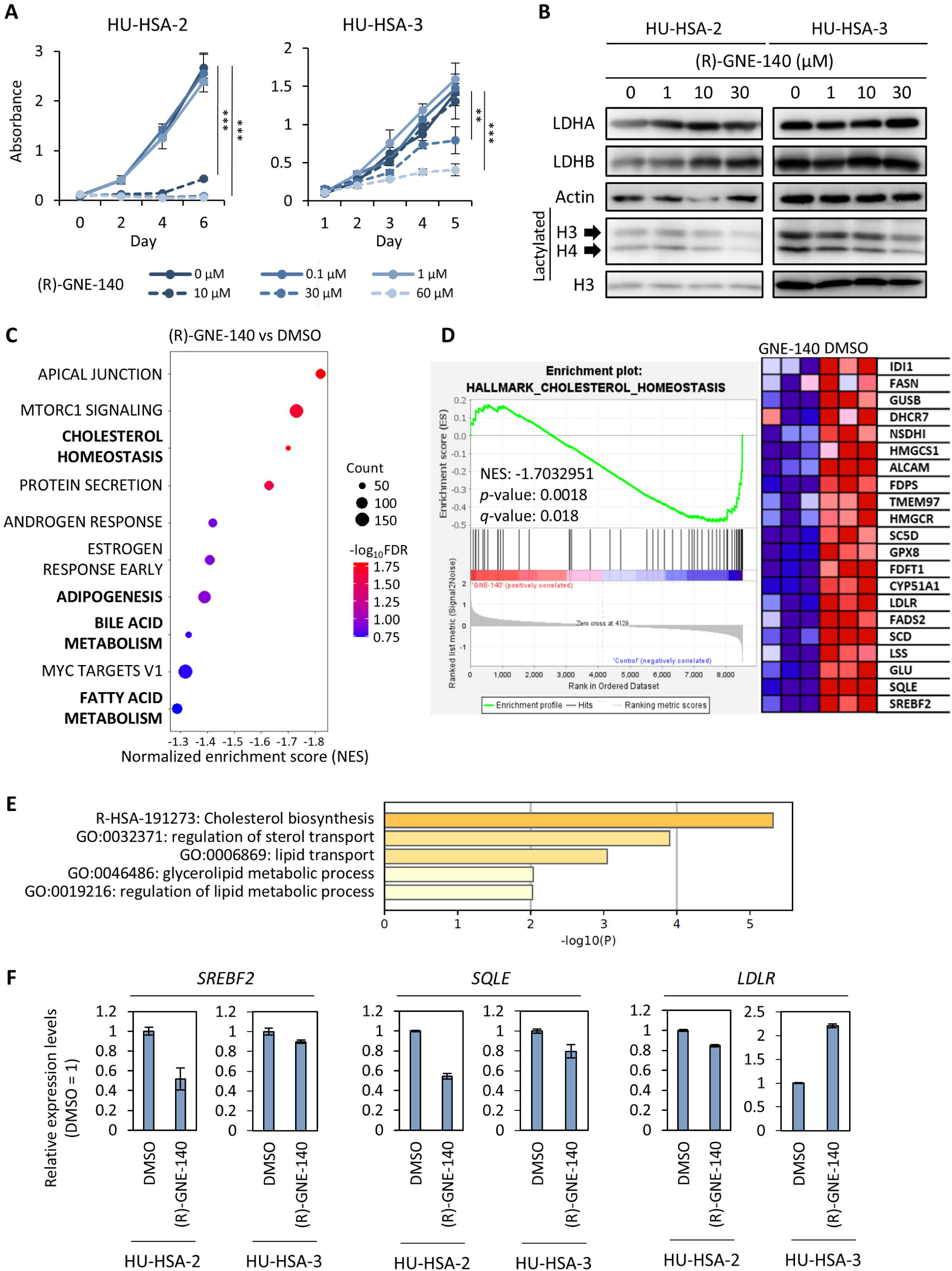
(R)-GNE-140 represses HSA cell proliferation and is associated with suppression of cholesterol/lipid metabolism in HU-HSA-2. **(A)** Cell proliferation curves of HU-HSA-2 and HU-HSA-3 treated with indicated concentrations of (R)-GNE-140. \*\**p* < 0.01, \*\*\**p* < 0.001. Dunnett’s test. Data are presented as average ± SD (*n* = 3). **(B)** Western blotting analysis for HU-HSA-2 and HU-HSA-3 cells 48 hours after treatment with indicated concentrations of (R)-GNE-140 (0 μM is the DMSO control). **(C-E)** GSEA and Metascape analyses of HU-HSA-2 cells treated with 10 µM (R)-GNE-140 for 72 hours compared to DMSO controls. (C) Dot plot of the top 10 negatively enriched pathways in GSEA. (D) GSEA enrichment plot (left) and heat map (right) of HALLMARK_CHOLESTEROL_HOMEOSTASIS gene set. (E) Gene Ontology enrichment analysis results generated using Metascape. **(F)** RT-qPCR analysis of representative cholesterol/lipid metabolism genes in HU-HSA-2 cells treated with 10 µM (R)-GNE-140 or DMSO for 72 hours and in HU-HSA-3 cells treated with DMSO or 30 µM (R)-GNE-140 for 48 hours. *B2M* and *TBP* were used in HU-HSA-2, and *RPL32* and *HMBS* were used in HU-HSA-3 as reference genes. Data are shown relative to DMSO controls (DMSO =1) and as average ± SD (*n* = 3).

Alternatively, sodium oxamate, a pyruvate analogue and competitive inhibitor of LDH, suppressed the proliferation of both HU-HSA-2 and HU-HSA-3 i*n vitro* in a dose-dependent manner (Fig. 2A). Although LDHA and LDHB protein levels were not altered, sodium oxamate decreased global histone lactylation levels in both cell lines (Fig. 2B). The expression levels of representative cholesterol/lipid metabolism genes were markedly reduced in HU-HSA-3, whereas changes in HU-HSA-2 were limited (Fig. 2C).

**Fig. 2.**
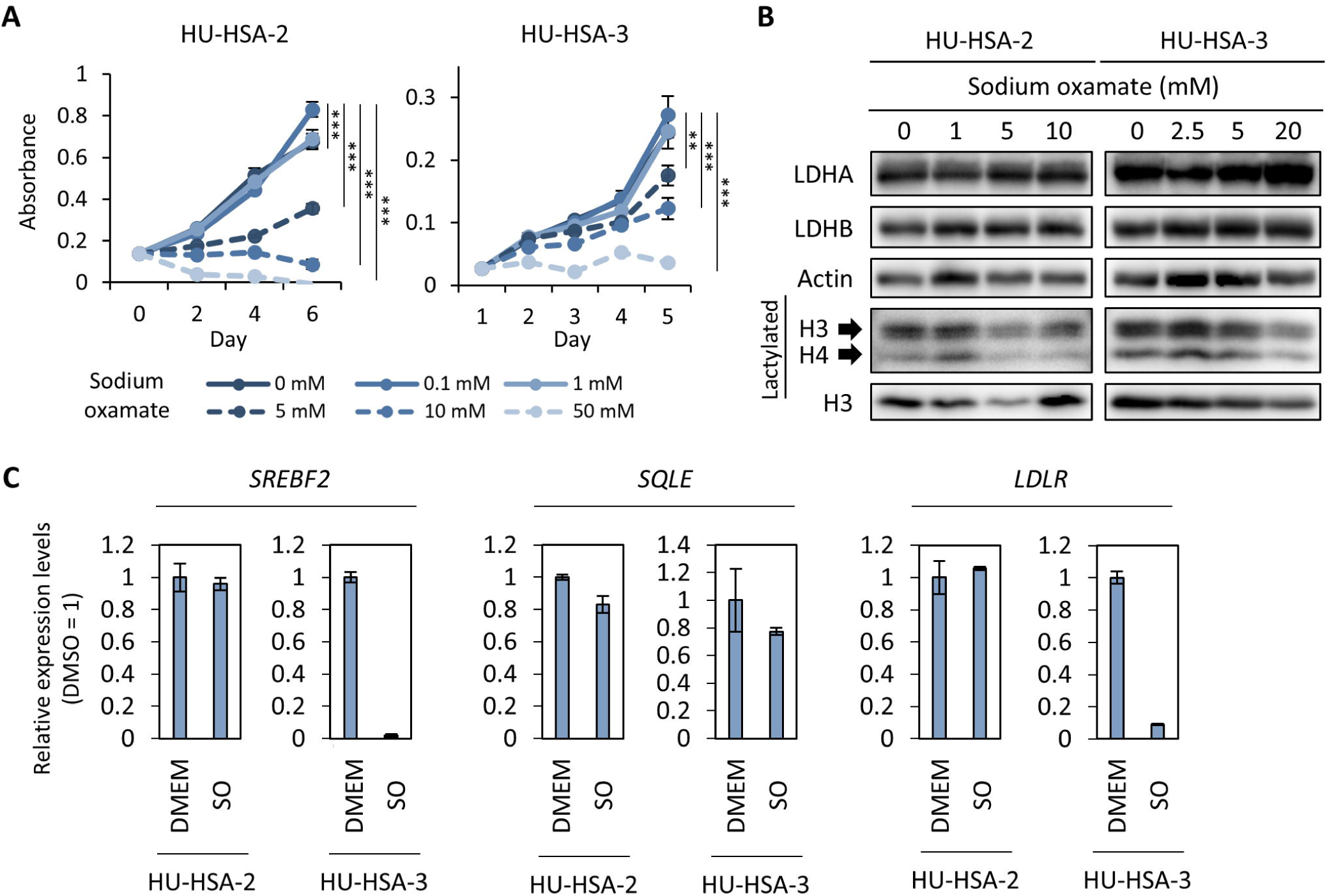
Sodium oxamate represses HSA cell proliferation and is associated with suppression of cholesterol/lipid metabolism in HU-HSA-3. **(A)** Cell proliferation curves of HU-HSA-2 and HU-HSA-3 treated with indicated concentrations of sodium oxamate. \*\**p* < 0.01, \*\*\**p* < 0.001. Dunnett’s test. Data are presented as average ± SD (*n* = 3). **(B)** Western blotting analysis for HU-HSA-2 and HU-HSA-3 cells 48 hours after treatment with indicated concentrations of sodium oxamate (0 mM is the DMEM control). **(C)** RT-qPCR analysis of representative cholesterol/lipid metabolism genes in HU-HSA-2 and HU-HSA-3 cells treated with 10 mM sodium oxamate (SO) or DMEM (control) for 48 hours. *RPL32* and *YWHAZ* were used as reference genes. Data are shown relative to DMEM controls (DMEM =1) and as average ± SD (*n* = 3).

To directly assess the contributions of LDHA and LDHB to cholesterol/lipid metabolism gene expression, we generated LDHA- and LDHB-knockout HU-HSA-3 clones (LDHA KO #1-4, LDHB KO #1-4, Fig. 3A). The expression levels of SREBP2 (*SREBF2*), a key regulator of cholesterol homeostasis, were reduced in both LDHA- and LDHB-knockout clones (Fig. 3A). Global histone lactylation levels were decreased in LDHA-knockout clones but were not consistently altered across LDHB-knockout clones (Fig. 3A). Cholesterol/lipid metabolism genes were broadly downregulated in LDHA-knockout clones, whereas LDHB-knockout clones showed variable effects (Fig. 3B). To assess the functional effects on intracellular lipid storage, we stained and quantified lipid droplets in HU-HSA-3 knockout clones. Both *LDHA* and *LDHB* knockout clones showed the reduced lipid droplet numbers compared to scramble controls (Fig. 4A - C).

**Fig. 3.**
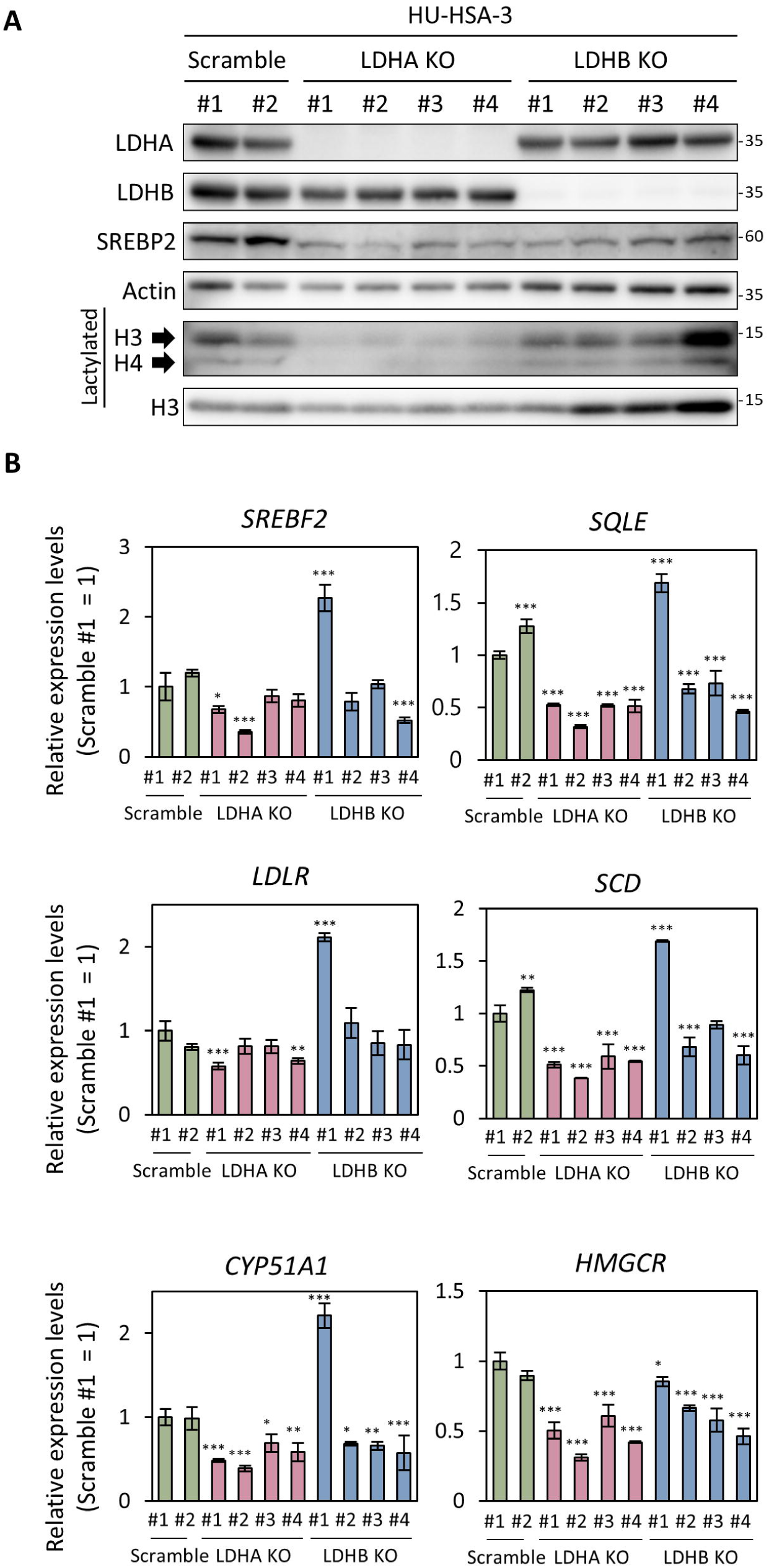
*LDHA* knockout was associated with downregulation of cholesterol/lipid metabolism genes. **(A)** Western blot analysis for scramble (control), *LDHA* knockout (LDHA KO), and *LDHB* knockout (LDHB KO) HU-HSA-3 clones. **(B)** RT-qPCR analysis of cholesterol metabolism/lipid metabolism genes in HU-HSA-3 clones. Gene expression was normalized to Scramble #1. Data are presented as average ± SD (*n* = 3). *HMBS* and *B2M* were used as reference genes. \**p* < 0.05, \*\**p* < 0.01, \*\*\**p* < 0.001. Dunnett’s test.

**Fig. 4.**
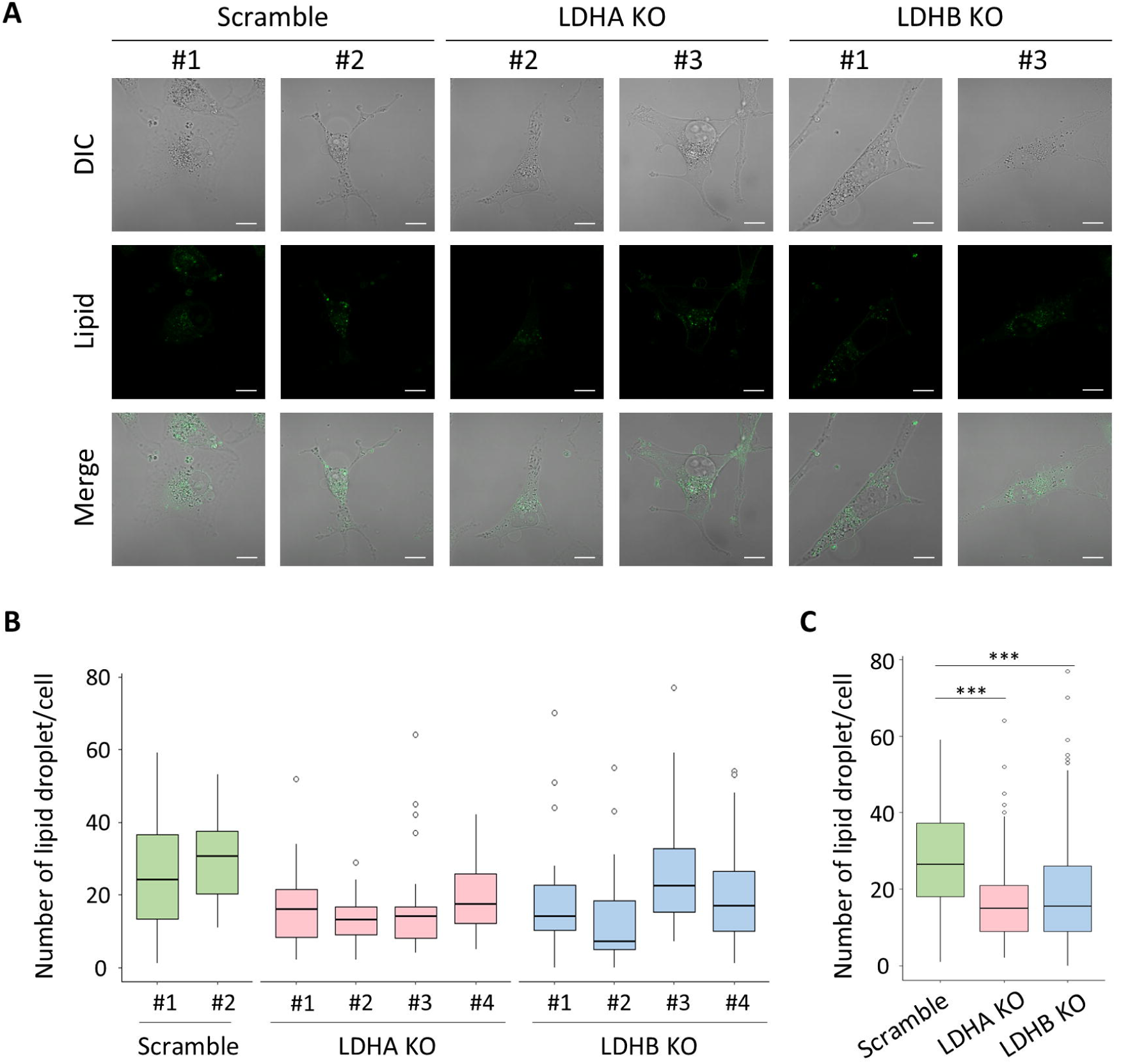
LDH knockout suppresses lipid droplet synthesis. **(A)** Representative images of HU-HSA-3 scramble control clones (#1 and #2), LDHA KO clones (#2 and #3), and LDHB KO clones (#1 and #3). Images show differential interference contrast (DIC), lipid droplet staining (Lipid), and merged images. Scale bars = 10 µm. **(B)** Box plots showing the number of lipid droplets per cell in individual clones (30 cells per clone). **(C)** Box plots showing the number of lipid droplets per cell in the pooled populations. (Scramble, *n* = 60; LDHA KO, *n* = 120; LDHB KO, *n* = 120). \*\*\**p* < 0.001. Kruskal-Wallis test followed by Dunn’s multiple comparisons test.

Together, these results suggest that LDH inhibition and genetic loss are associated with changes in cholesterol/lipid metabolism gene expression, and that responsiveness to LDH inhibitors differs between HU-HSA-2 and HU-HSA-3 cell lines.

### Statins inhibit HU-HSA cell proliferation *in vitro*, and fluvastatin plus dipyridamole suppresses HSA PDX tumor growth

Given the observed association between LDH perturbation and cholesterol/lipid metabolism, together with reduced SREBP2 (*SREBF2*) levels, we assessed whether inhibition of the cholesterol biosynthesis pathway with statins suppresses HSA cell growth. Fluvastatin and rosuvastatin, HMG-CoA reductase inhibitors, significantly reduced cell growth of both HU-HSA-2 and HU-HSA-3 cells *in vitro* (Fig. 5A, B). Fluvastatin increased cleaved caspase-3 levels, an apoptosis marker, in both cell lines, and it reduced phospho-histone H3 (pH3S10) levels in HU-HSA-3 (Fig. 5C). To assess its anti-tumor activity *in vivo*, we treated canine HSA PDX models (HU-HSAPDX-3 and HU-HSAPDX-5) with fluvastatin intraperitoneally; however, no statistically significant suppression of tumor growth was observed in either model (Fig. 5D).

**Fig. 5.**
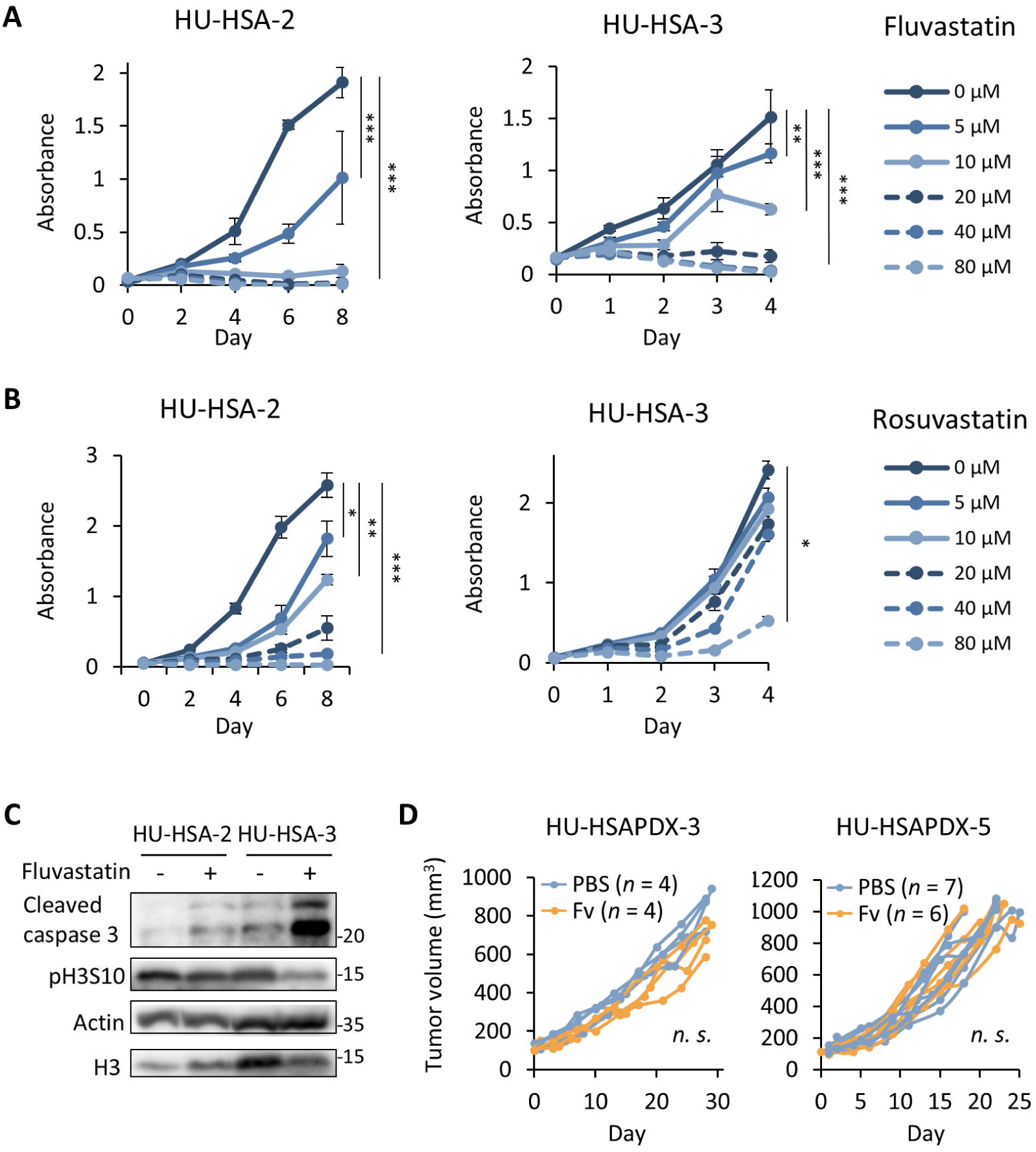
Statins inhibit HSA cell proliferation *in vitro*. **(A, B)** Cell proliferation curves of HU-HSA-2 and HU-HSA-3 treated with indicated concentrations of fluvastatin (A) or rosuvastatin (B). \**p* < 0.05, \*\**p* < 0.01, *** *p* < 0.001. Dunnett’s test. Data are presented as mean ± SD (*n* = 3). **(C)** Western blot analysis for HU-HSA-2 and HU-HSA-3 treated with DMSO or 10 µM fluvastatin for 48 hours. **(D)** Tumor growth curves of HU-HSAPDX-3 and HU-HSAPDX-5 treated with PBS or fluvastatin (Fv) intraperitoneally. *n.s.*, not significant. Two-way ANOVA.

To enhance the inhibitory effect on the cholesterol biosynthesis pathway, we evaluated fluvastatin in combination with dipyridamole. Dipyridamole reduced HU-HSA-2 and HU-HSA-3 cell viability *in vitro* in the micromolar range (Fig. 6A), and co-treatment with fluvastatin produced overall additive effects in both cell lines (ZIP synergy score: 7.656 ± 1.2 for HU-HSA-2 and 4.236 ± 0.96 for HU-HSA-3; Fig. 6B). Finally, we treated canine HSA PDX models (HU-HSAPDX-6 and HU-HSAPDX-7) with fluvastatin (p.o.) and/or dipyridamole (i.p.). While dipyridamole monotherapy showed modest tumor growth inhibition, combination treatment resulted in greater tumor growth reduction in both PDX models (Fig. 6C). Body weight remained stable across treatment groups (Fig. 6D).

**Fig. 6.**
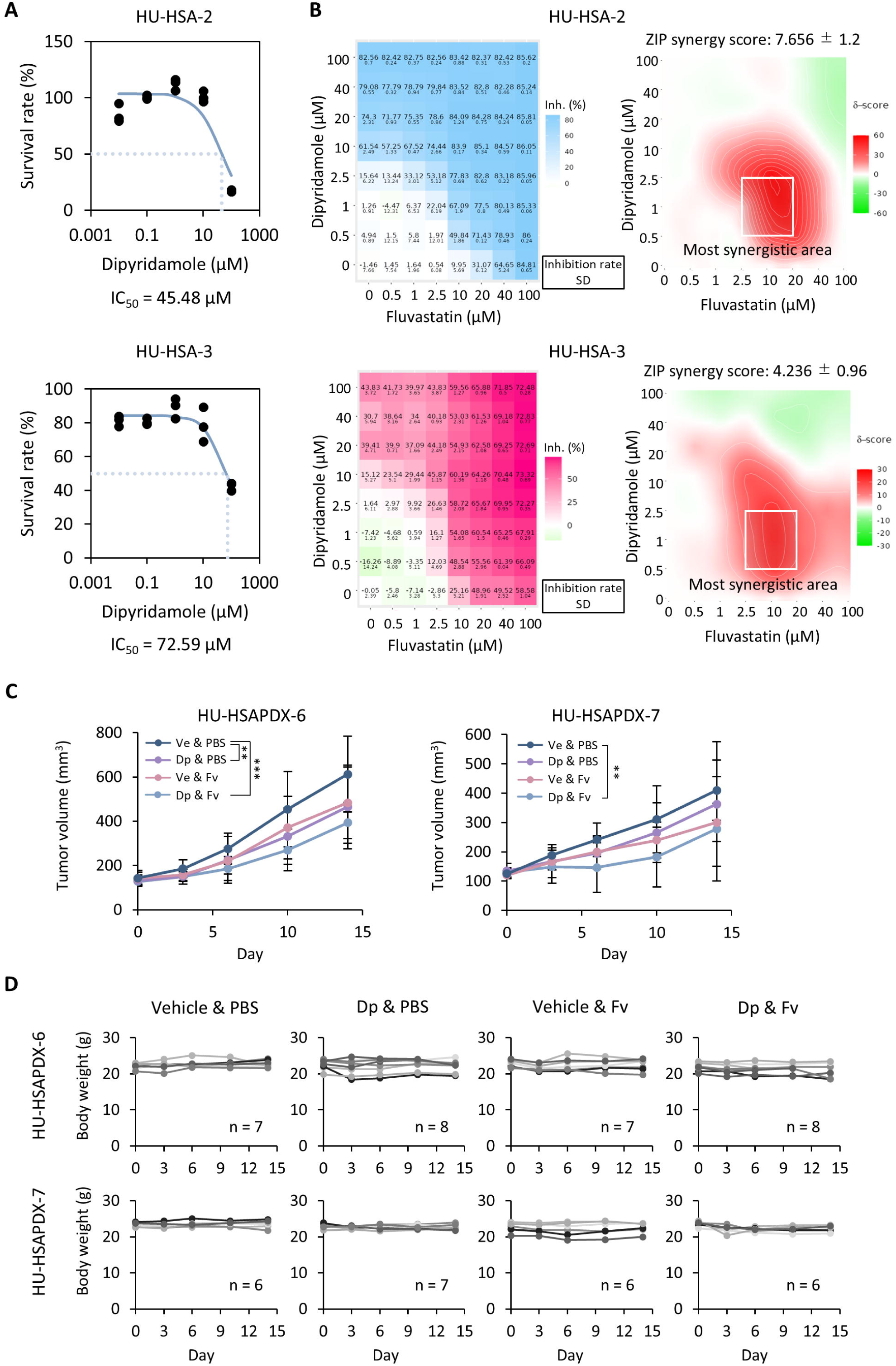
Co-treatment with fluvastatin and dipyridamole suppresses PDX tumor growth. **(A)** Survival curves and IC_50_ values for HU-HSA-2 (top) and HU-HSA-3 (bottom) cells treated with dipyridamole for 48 hours. **(B)** Inhibition matrices (left) and synergy maps (right) for HU-HSA-2 (top) and HU-HSA-3 (bottom) cells treated with dipyridamole and fluvastatin. Values in the inhibition matrices are presented as mean ± SD (*n* = 3). Synergy was evaluated using the ZIP model. ZIP synergy scores are shown. The most synergistic area is boxed. **(C)** Tumor growth curves of HU-HSAPDX-6 and HU-HSAPDX-7 treated with vehicle (Ve) or dipyridamole (Dp) in combination with PBS or fluvastatin (Fv). \*\**p* < 0.01, *** *p* < 0.001. Two-way ANOVA. Data are presented as mean ± SD (*n* = 6–8 per group). **(D)** Body weight plots of individual mice in the co-treatment experiments for HU-HSAPDX-6 and HU-HSAPDX-7.

These results indicate that although fluvastatin monotherapy showed limited activity *in vivo*, co-treatment with dipyridamole could enhance tumor growth inhibition in PDX models.

## Discussion

This study supports a functional association between LDH and cholesterol/lipid metabolism in HSA cells and suggests that this axis may be therapeutically exploitable when the sterol-regulated feedback loop is concurrently inhibited.

Our data indicate that the relationship between LDH perturbation and cholesterol/lipid metabolism is context dependent. In HU-HSA-2, transcriptome profiling after (R)-GNE-140 treatment identified cholesterol/lipid metabolism gene sets among the most negatively enriched pathways, whereas (R)-GNE-140 did not comparably suppress representative cholesterol/lipid metabolism genes in HU-HSA-3. Conversely, sodium oxamate reduced cholesterol/lipid metabolism gene expression in HU-HSA-3 but not HU-HSA-2. This difference may reflect biological heterogeneity at both the inter□tumor level (patient□to□patient differences) and the intra□tumor level (distinct subpopulations within a tumor tissue).^25^ Our knockout experiments suggest that LDHA plays an important role in regulating cholesterol/lipid metabolism gene expression in HU-HSA-3. *LDHA* knockout clones showed a consistent reduction in expressions of these genes, whereas *LDHB* knockout clones showed variable effects. In parallel, global histone lactylation decreased robustly in LDHA knockout clones but was not consistently altered across LDHB knockout clones. These findings suggest that cholesterol/lipid metabolism gene regulation is more tightly coupled to LDHA than to LDHB alone, potentially reflecting isoform-specific substrate preferences and compartmentalization (e.g., nuclear LDHA and mitochondrial-associated LDHB in some contexts).^2,3,26^ However, *LDHB* knockout clones also showed reduced SREBP2 protein levels and decreased lipid droplet numbers. This suggests that LDHB also contributes to cholesterol biosynthesis, possibly through mechanisms different from LDHA. To define how broadly this LDH-cholesterol/lipid metabolism relationship applies to HSA pathogenesis and to fully understand its mechanism, additional HSA models such as more primary cell lines, cell line-derived xenografts, and PDX models, should be studied.

Mechanistically, LDH inhibition may modulate cholesterol/lipid metabolism through coupled redox control and lactate-linked chromatin regulation. The LDH reaction couples pyruvate-lactate interconversion to the regeneration of cytosolic NAD□ from NADH.^27,28^ LDH inhibition can promote reductive stress (an increased cytosolic NADH/NAD□ ratio), which can constrain glycolytic flux and downstream acetyl-CoA production.^27,28^ These metabolic changes potentially attenuate nutrient-sensing pathways such as the mTORC1 pathway, which in turn suppresses SREBP activation and the subsequent transcriptional program for cholesterol biosynthesis.^9,29^ Consistent with this, our HU-HSA-2 transcriptome analysis showed coordinated downregulation of cholesterol/lipid gene sets together with reduced mTORC1 signaling following (R)-GNE-140 exposure. In parallel, reduced intracellular lactate availability may limit the substrate for histone lactylation.^4,6^ The concurrent reduction in global histone lactylation and lipid droplet accumulation observed in LDHA-knockout HU-HSA-3 cells suggests the possibility that histone lactylation contributes to the transcriptional maintenance of cholesterol/lipid pathways. Establishing a direct causal link, however, requires demonstrating that lactylation marks are specifically enriched at the promoters or other regulatory regions of these metabolic genes.

The inhibitory effects on LDH exerted by small-molecule inhibitors and genetic knockout should be interpreted with caution. (R)-GNE-140 is a potent small-molecule LDH inhibitor whose binding is reported to be NADH-dependent, likely reflecting preferential engagement of the NADH-bound enzyme state.^30^ By contrast, sodium oxamate is a pyruvate analogue that competitively inhibits the substrate-binding site on the LDH-NADH complex.^31^ Given that sodium oxamate competes with intracellular pyruvate, its efficacy is expected to vary according to pyruvate abundance and the broader metabolic state. Specifically, HU-HSA-3 cells rely more heavily on glucose for ATP production than HU-HSA-2.^6^ Consequently, sodium oxamate might perturb pyruvate metabolism and redox homeostasis beyond simple LDH inhibition, potentially contributing to the observed decrease in cholesterol/lipid gene expression in HU-HSA-3 cells.^31,32^ Furthermore, knocking out either LDHA or LDHB can induce compensatory remodeling of LDH tetramers, whereas small-molecule inhibitors suppress the activity of all LDH tetramers regardless of their subunit composition.^33,34^ Collectively, while these factors have to be carefully taken into account, our data indicate an association between LDH and cholesterol/lipid metabolism.

The enhanced antitumor activity of the fluvastatin-dipyridamole combination in our PDX models is consistent with prior human cancer studies.^10,11^ Statin-mediated inhibition of HMG□CoA reductase can activate an SREBP2□driven sterol□regulated feedback response that limits statin cytotoxicity, whereas dipyridamole attenuates this response.^10,11^ In both co□treated HSA PDX models (HU□HSAPDX□6 and HU□HSAPDX□7), tumor growth was significantly delayed compared with either fluvastatin or dipyridamole monotherapy; however, co-treatment did not induce overt tumor regression. This suggests that fluvastatin-dipyridamole co-treatment might be insufficient to shrink established tumors at least under the tested condition. Nonetheless, a cytostatic effect may still have clinical utility in canine HSA, in which metastasis often occurs prior to or shortly after diagnosis, and systemic therapy is commonly conducted following surgical removal of primary tumors.^35,36^ Although doxorubicin-based adjuvant therapy is associated with longer survival than surgery alone, overall outcomes remain poor.^37^ Accordingly, additional systemic strategies that delay metastatic/disseminating progression may improve patient prognosis. Therefore, it may be more clinically relevant to evaluate the efficacy of fluvastatin-dipyridamole co-treatment in metastatic or disseminating conditions. To this end, the development of HSA models such as cell-line derived xenografts and PDX models capable of metastasis or dissemination is essential for future preclinical evaluations.

In conclusion, we identified a functional link between LDH and cholesterol/lipid metabolism in canine HSA cell lines and demonstrated that fluvastatin-dipyridamole co□treatment significantly delayed tumor growth in HSA PDX models. Further studies are warranted to elucidate the mechanisms by which LDH regulates transcription of genes involved in cholesterol/lipid metabolism and to define the clinical potential of combined statin-dipyridamole therapy.

## Acknowledgments

We thank Dr Hironobu Yasui (Faculty of Veterinary Medicine, Hokkaido University) for helpful advice and constructive discussions. We are grateful to Drs Kenji Hosoya, Sangho Kim, and Ryohei Kinoshita (Veterinary Teaching Hospital, Faculty of Veterinary Medicine, Hokkaido University) for providing patient samples, and to all members of the Laboratory of Comparative Pathology (Faculty of Veterinary Medicine, Hokkaido University) for helpful discussions, encouragement, and support. We sincerely thank the canine patients whose tumor tissues were used to generate cell lines and patient-derived xenograft models for this study, and we are deeply grateful to their owners for their generous cooperation in supporting this research.

## Author contributions

T.S., S.T., and K.A. designed the research. K.A. conceived and supervised the project. T.S. and S.T. performed most experiments. T.S., S.T., K.K., T.G., and K.A. performed the dual-drug treatment experiments. K.A. conducted the bioinformatic analyses. T.S. and K.A. acquired funding. T.S., S.T., and K.A. wrote the original draft. All authors reviewed and edited the manuscript.

## Use of AI-assisted tools

We used OpenAI ChatGPT (model 5.2 Pro) and Google Gemini 2.5 Pro for English proofreading and methodological suggestions. All outputs were reviewed and edited by the authors. No AI-generated data were used.

## Data and materials availability

mRNA-seq data have been deposited in the Gene Expression Omnibus (GEO) under accession GSE314239. During peer review, reviewers may access the private record using the reviewer token kxanmyocrbetbeh at https://www.ncbi.nlm.nih.gov/geo/query/acc.cgi?acc=GSE314239. These data will be made publicly accessible upon publication. All non-commercially available new materials, including constructs, cell lines, and PDX models, that Hokkaido University has the right to provide will be made available to non-profit or academic requesters upon completion of a standard material transfer agreement. Requests for materials should be addressed to K.A.

