## Supplemental information for "Lactate dehydrogenase is associated with cholesterol/lipid metabolism, and fluvastatin plus dipyridamole suppresses canine hemangiosarcoma growth in patient-derived xenograft models"

The PDF file includes:

Materials and Methods

Supplementary Figure

Supplementary Tables

References

#### Materials and Methods

##### *Cell culture*

Two canine HSA cell lines (HU-HSA-2 and HU-HSA-3), isolated from splenic hemangiosarcoma canine patients,[1] and 293T cells, obtained from RIKEN Bioresource Center, were cultured in Dulbecco's Modified Eagle Medium (DMEM; #044-29765, FUJIFILM Wako Pure Chemical Corporation, Osaka, Japan) supplemented with 10% fetal bovine serum (FBS; #10270-106, Thermo Fisher Scientific, Waltham, MA, USA) and penicillin-streptomycin (P/S; #168-23191, FUJIFILM Wako) at 37°C with 5% $CO_2$. For HU-HSA-2 and HU-HSA-3 cells, cell culture dishes were precoated with 0.1% gelatin derived from bovine bone (#074-02761, FUJIFILM Wako) by incubation for 30 minutes at 37°C.

##### *Cell line authentication statement*

No additional authentication was performed in this study. HU-HSA-2 and HU-HSA-3 cells were established and authenticated in a previous study.[1] 293T cells were authenticated in RIKEN bioresource center (RCB2202).

##### *LDHA and LDHB knockout cell generation*

293T cells were cultured in CELLSTAR® 6-well cell culture multiwell plates (#65710, Greiner Bio-One, Kremsmünster, Austria) until they reached 70% confluency. The next day, medium was replaced with DMEM containing 10% FBS without P/S, and cells were transfected with 4 µg lentiCRISPRv2 containing designed single guide RNAs (sgRNAs), 0.5 µg HIV-gp, and 0.5 µg VSV-G using 8.25 µL Lipofectamine 3000 with 7.5 µL P3000 reagent (#L300015 Thermo Fisher Scientific) in 250 µL Opti-MEM (#31-985-070, Thermo Fisher Scientific) per well according to the manufacturer's instructions. Forty-eight hours after transfection, medium was filtered using Minisart 0.45 µm filters

(#S6555-FMHUK, Sartorius, Göttingen, Germany) and collected in 15 mL tubes as viral solutions. Polybrene (#12996-81, Nacalai Tesque, Kyoto, Japan) was added to the viral solutions to a final concentration of 10 μg/mL. Fifty thousand HU-HSA-3 cells were seeded with virus solution into 6-well plates. Eight hours after infection, the viral solution was removed and replaced with fresh culture medium. Cells were cultured for 2-3 days until they reached 60%-80% confluency, and medium was replaced with medium containing 4 μg/mL puromycin. Once cells started stably proliferating in the selection medium, they were seeded into 96-well plates at a density of 0.5 cells per well. When the cells reached 70%–90% confluency, they were passaged to larger-scale plates. Total protein was harvested from 12-well culture plates, and knockout efficiency was evaluated by western blotting.

*Cell proliferation assay*

Cells were seeded in CELLSTAR® 12-well cell culture multiwell plates (#665180, Greiner Bio-One) with 1 mL culture medium or 96-well plates (#655180, Greiner Bio-One) with 100 μL culture medium. After cells were attached, they were treated with sodium oxamate (#327-24621, FUJIFILM Wako), (R)-GNE-140 (#S6675, Selleck Biotechnology, Yokohama, Japan), fluvastatin (#068-06641, FUJIFILM Wako), rosuvastatin (#187-03361, FUJIFILM Wako), or dipyridamole (#D9766, Sigma-Aldrich, St. Louis, MO, USA). DMEM or dimethyl sulfoxide (DMSO) were used as controls for each compound. The number of live cells was counted using Countess II (AMQAX1000, Thermo) after trypan blue staining or using Cell Counting Kit-8 (CCK-8; #343-07623, Dojindo Laboratories, Kumamoto, Japan) according to the manufacturer's instructions. Relative viability was calculated by setting control samples as 100%. KyPlot 6.0 software (KyensLab Inc., Tokyo, Japan) was used to generate survival curves.[2] For cell proliferation curves, cells were subjected to CCK-8 assays every other day (HU-HSA-2) or every day (HU-HSA-3) according to the manufacturer's instructions. For drug combination analysis, HU-HSA-2 and HU-HSA-3 cells seeded in 96-well plates were treated for 48 hours with fluvastatin and dipyridamole in an 8 × 8 dose–response matrix (0, 0.5, 1, 2.5, 10, 20, 40, and 100 μM for each drug). Cell viability was assessed using the CCK-8 assay and normalized to DMSO-treated controls, then converted to percent inhibition (100 - % viability). Drug interaction was quantified using SynergyFinder 3.0 with the Zero Interaction Potency (ZIP) reference model.[3,4] ZIP δ-scores were calculated for each dose pair as the difference between the observed inhibition and the ZIP-predicted inhibition, and the overall ZIP synergy score was reported as the mean δ-score across the entire matrix (mean ± SD, n = 3). ZIP synergy scores were interpreted as synergistic (>10), additive (−10 to 10), or antagonistic (<−10). The most synergistic 3 × 3 dose window (most synergistic area) was highlighted on the synergy maps.

*Protein extraction and western blotting*

SDS lysis buffer {2% SDS, 50 mM Tris-HCl (pH6.8), 1 mM EDTA (pH8.0)} was added to cultured cells after washing with phosphate-buffered saline (PBS) twice. Cell lysates were sonicated using BRANSON Sonifier 450 (Branson Ultrasonics Corporation, Danbury, CT, USA) for 2 seconds at power 2. Protein concentrations were measured with TaKaRa BCA Protein Assay Kit (#T9300A, Takara Bio Inc., Kusatsu, Shiga, Japan) before adding 4× sample loading buffer {200 mM Tris-HCl buffer (pH 6.8), 8% SDS, 40% glycerol, 1% bromophenol blue, 20% 2-mercaptoethanol} and denaturing at 98°C for 5 minutes. One to 3 μg of protein were separated on 8%-18% SDS-polyacrylamide gels by electrophoresis and transferred to Immobilon-P transfer membranes (#IPVH00010, Merck Millipore, Burlington, MA, USA). Membranes were blocked with 3% skim milk in Tris-buffered saline with 0.05% Tween20 (TBST) for 1 hour at room temperature (RT) and incubated with primary antibodies in Can Get Signal Solution 1 (#NKB-101, TOYOBO Co., Ltd., Osaka, Japan) overnight at 4°C. After washing with TBST three times, membranes were incubated with the corresponding secondary anti-mouse (#G21040) or anti-rabbit (#G21234) IgG antibody (Thermo Fisher Scientific) in 3% skim milk for 1 hour at RT. Signals were developed with Immobilon Western Chemiluminescent HRP substrate (#WBKIS0500, Merck Millipore) and visualized using an ImageQuant LAS 4000 mini luminescent image analyzer (Cytiva, Marlborough,

MA, USA). Data were processed using ImageJ software (v. 1.54p).[5] Antibodies used in this study are listed in Table S1.

*Generation of anti-canine LDHA antiserum*

A custom rabbit polyclonal antiserum against canine LDHA was generated by Cosmo Bio Co., Ltd. (Tokyo, Japan). A synthetic peptide corresponding to a canine LDHA-specific sequence (ISDVVKVTLTPEEEARLKK) was designed to minimize cross-reactivity with canine LDHB. Rabbits were immunized with the peptide antigen according to the following schedule: primary immunization on day 0, followed by booster immunizations on days 14, 28, and 42. Whole blood was collected on day 49, and the antiserum was used for experiments without further purification. Antibody titers were assessed by ELISA and confirmed to exceed the manufacturer's internal quality-control threshold.

*mRNA-seq*

HU-HSA-2 cells were treated with DMSO or 10 μM (R)-GNE-140 for 72 hours in triplicate. Total RNA was extracted with NucleoSpin RNA isolation kit (#740955.50, Macherey-Nagel GmbH & Co. KG, Düren, Germany) according to the manufacturer's instructions, and submitted to Rhelixa (Tokyo, Japan) for library preparation and sequencing. Libraries were generated with NEBNext Poly(A) mRNA Magnetic Isolation Module (E7490, New England Biolabs, Ipswich, MA, USA) and NEBNext Ultra II Directional RNA Library Prep Kit (E7760, New England Biolabs). Sequencing was conducted with NovaSeq 6000 (Illumina, San Diego, CA, USA). Sequence reads were mapped to ROS_Cfam_1.0 (CanFam 4) using STAR,[6] and expression levels were estimated using RSEM.[7] Differential expression analysis was performed using edgeR, and gene expression profiles were analyzed by gene set enrichment analysis (GSEA) v4.1.0.[8,9] Gene Ontology analysis was conducted using Metascape.[10]

Sequence data and processed files were uploaded to the Gene Expression Omnibus (GSE314239).

*RNA extraction and reverse transcription quantitative polymerase chain reaction (RT-qPCR)*

Total RNA was extracted by using Tripure Isolation Reagent (#11667157001, Roche, Basel, Switzerland), and reverse transcription was performed using PrimeScript Reverse Transcriptase II (#2690A, Takara Bio) according to the manufacturer's instructions. One microgram of total RNA was used for reverse transcription to obtain cDNA. qPCR was performed using the cDNAs and primers listed in Table S2. Sample preparation was performed using KAPA SYBR Fast qPCR Kit Master Mix (2×) ABI Prism (#KK4604, KAPA Biosystems, Wilmington, MA, USA). All samples were applied in triplicate and analyzed with StepOne Real Time PCR System (Thermo Fisher Scientific). Samples were denatured at 95°C for 10 minutes, followed by 40 cycles of 95°C for 3 seconds and 60°C for 30 seconds. Results were normalized based on the geometric mean of reference genes, which were selected from seven potential internal controls (*RPL32, RPL13A, TBP, YWHAZ, HMBS, B2M,* and *ACTB*) using geNorm software.[11] Selected reference genes for each experiment are listed in Table S3. Ensembl and Primer3 software were used to design primers that can target all splicing variants and exon-exon junctions. The BLAST database was used to confirm that each primer sequence did not detect other genes.[12] Primer set specificity was evaluated by confirming that each primer set had an identical and singular peak in the melting curve. Relative expression levels were calculated by setting the corresponding control samples to 1.0.

*Lipid droplet staining*

For visualization of lipid droplets, scramble, LDHA, and LDHB knockout HU-HSA-3 clones were seeded at approximately 70% confluency in gelatin-coated 35-mm glass-bottom dishes (#D11130H, Matsunami Glass Ind. Ltd., Osaka, Japan) and incubated overnight. After washing twice with PBS, cells were fixed with 4% paraformaldehyde for 7.5 minutes at RT. Cells were then washed twice with PBS and stained with Lipi-Green (#LD02, Dojindo) diluted in PBS according to the manufacturer's instructions, followed by incubation for 30 minutes at 37 °C. After a final PBS wash, images were acquired using a confocal laser microscope (LSM 800, Carl Zeiss, Oberkochen, Germany) with a ×63 objective. For quantification of lipid droplets, 30 confocal images (one cell per image) were analyzed for each clone and quantified using ImageJ.[5] Individual cells were manually outlined as regions of interest. Images were converted to 8-bit and thresholded using Li's method (dark background).[13] A binary mask was generated, droplets were separated using the watershed algorithm, and particles were quantified with Analyze Particles (size: 0.0005-0.10; circularity: 0-1.00).

*Plasmid construction*

*LDHA* (ENSCAFG00000009211) and *LDHB* (ENSCAFG00000012195) sequences were obtained from Ensembl, and sgRNA sequences for each gene were designed using CHOPCHOP.[14] Scramble sgRNA sequences were obtained by randomizing sgLDHB sequences and were confirmed to have no highly matched sequences in the canine genome using BLAST search.[12] sgRNA sequences used in this study are listed in Table S4. sgRNA oligos were synthesized by Eurofins Genomics (Tokyo, Japan) and used for plasmid construction. Forward and reverse oligos for each target gene were mixed with T4 polynucleotide kinase (#2021S, Takara Bio) and annealed using the following program: 37°C for 30 minutes, 95°C for 5 minutes, then ramp-down to 25°C at a 4.1% rate, using Veriti 96-Well Fast Thermal Cycler (Thermo Fisher Scientific). Annealed oligos were ligated to lentiCRISPRv2 (a gift from Feng Zhang; Addgene plasmid # 52961; RRID: Addgene_52961) linearized by BsmBI-v2 digestion (R0739S, New England Biolabs), using DNA Ligation Mix (#2011A, Takara Bio) at 16°C overnight. A total of 2.5 µL ligation product was added to 25 µL NEB Stable competent *E. coli* (#3040, New England Biolabs) and incubated on ice for 30 minutes. Samples were heated at 42°C for 45 seconds, followed by incubation on ice for 2 minutes. After adding 100 µL SOC medium, competent cell solutions were spread on LB plates containing 100 µg/mL ampicillin and incubated at 37°C overnight. A single colony was picked and cultured in 2 mL Terrific Broth medium containing 100 μg/mL ampicillin at 37°C for 24 hours. Plasmids were extracted with NucleoSpin plasmids EasyPure (#U0727, Macherey-Nagel GmbH & Co. KG,) according to the manufacturer's instructions.

*Establishment and characterization of canine PDX models*

Canine HSA PDX models were established from fresh hemangiosarcoma tissues obtained from canine patients that underwent splenectomy with written informed consent from the owners and approval from the Ethics Screening Committee of an animal hospital (2022-005). All cases were confirmed as hemangiosarcoma by two board-certified veterinary pathologists. Patient information is provided in Table S5. Tumor tissues were used for PDX development immediately after surgical resection. They were fragmented into approximately 2-3 mm cubes and then subcutaneously transplanted into both flanks of 6- to 8-week-old male or female KSN/Slc mice (Japan SLC, Inc. Shizuoka, Japan) according to the animal study protocol described below. Meloxicam (0.2 mg/kg) was intraperitoneally injected for analgesia on the day of surgery and again the following day. Tumors were resected when the volume reached 1 $cm^3$, re-fragmented, and then transplanted into other mice. This passaging procedure was repeated three times, after which the tissues were designated HU-HSAPDX-3, HU-HSAPDX-5, HU-HSAPDX-6, and HU-HSAPDX-7, and used as canine HSA PDX models. Characterization and authentication of HU-HSAPDX-3 were performed previously.[1] For HU-HSAPDX-5, HU-HSAPDX-6, and HU-HSAPDX-7, histopathological examination and immunohistochemistry for the endothelial markers CD31 and von Willebrand factor (vWF) were performed to confirm that the PDX models retained features of the original patient tumors. These three PDX models were authenticated by short tandem repeat (STR) analysis using the Canine Genotypes Panel 2.1 Kit (F864S, Thermo Fisher Scientific) (Fig. S1). Fragment analyses were conducted by Fasmac (Kanagawa, Japan).

*Hematoxylin and eosin (HE) staining, and immunohistochemistry (IHC)*

Tumor samples from PDX models were fixed in 10% neutral-buffered formalin, dehydrated through an ethanol series, cleared with xylene and infiltrated with paraffin wax in Tissue-Tek VIP5 Jr (Sakura Finetek Japan Co.,Ltd., Tokyo, Japan). The samples were embedded in paraffin wax and sliced into 2 μm-thick sections. For hematoxylin and eosin staining, tissues were deparaffinized with xylene and placed in 99%, 95%, 90%, 80%, and 70% ethanol for 2 minutes each in this order. After washing out the ethanol with tap water and distilled water (DW), the tissues were stained with hematoxylin for 1 minute and then washed with tap water for 5 minutes. Then, they were stained with eosin for 1.5 minutes after being placed in 95% ethanol for 2 minutes. Remaining eosin was washed with 95% ethanol. Afterwards, the tissues were dehydrated with absolute ethanol and cleared with xylene. Finally, the tissues were mounted with Eukitt (#6.00.01.0001.06.01.EN, ORSAtec GmbH, Bobingen, Germany) and covered with coverslips for histopathological analysis. For IHC, after deparaffinization, the tissues were washed with PBS three times, and antigens were retrieved in 1 mM EDTA in an autoclave machine for 5 minutes for CD31, and 20 μg/mL proteinase K in Tris-EDTA pH 8.0 at 37°C for 15 minutes for vWF. Endogenous peroxidases were inactivated with 0.3% $H_2O_2$ in methanol for 25 minutes at RT before blocking the tissue sections with 10% normal goat serum (#426042, Nichirei Biosciences Inc., Tokyo, Japan) for 30 minutes at RT. The sections were then incubated with anti-CD31 or vWF antibody overnight at 4°C. Afterwards, the tissues were washed with PBS three times and stained with peroxidase-conjugated goat anti-mouse IgG antibody (#424132, Nichirei Biosciences) or goat anti-rabbit IgG antibody (#424142, Nichirei Biosciences) for 30 minutes at RT. The slides were washed with PBS three times again, and signals were developed by reaction with 3,3'-diaminobenzidine (#349-00903, Dojindo). Antibodies used in this study are listed in Table S1.

*Animal studies*

All mouse experiments were performed under the guidelines (protocol number: 20-0083, 21-0062, 25-0062). Female 4- to 8- week-old KSN/Slc mice (Japan SLC, Inc.) were used for experiments. Mice were bred and raised at the animal facility under SPF conditions. Tumor fragments (3-mm cubes) of HU-HSAPDX-3, HU-HSAPDX-5, HU-HSAPDX-6, or HU-HSAPDX-7 were transplanted subcutaneously in the right flank of mice. Tumor volumes were calculated using the formula: volume = (length × width$^2$)/2. When the tumor volume reached 100 mm$^3$, treatments were started. For HU-HSAPDX-3 and HU-HSAPDX-5, mice were intraperitoneally injected with fluvastatin (30 mg/kg) every other day. PBS was used as the vehicle control. The treatment duration was up to 30 days or until the tumor volume reached 1 cm$^3$. For HU-HSAPDX-6 and HU-HSAPDX-7, mice were orally administered with fluvastatin (50 mg/kg) and/or injected intraperitoneally with dipyridamole (120 mg/kg) daily. Dipyridamole was dissolved in a solvent of 50% (v/v) polyethylene glycol 600 and 2 mg/mL tartaric acid to achieve a final concentration of 5 mg/mL, and this solvent was also used as the vehicle control for dipyridamole. PBS was used as the vehicle control for fluvastatin. The treatment duration was 2 weeks. Mice were euthanized with $CO_2$ at the end of treatments.

*Statistical analysis*

Statistical analyses were performed with Microsoft Excel (version 2019) and R software (version 4.2.0). For *in vivo* experiments, tumor growth was analyzed with two-way ANOVA. Cell proliferation was analyzed with two-way ANOVA and Dunnett's test. Statistical comparisons for lipid droplet quantification and qPCR among the three groups (scramble, LDHA knockout, and LDHB knockout clones) were performed using the Kruskal-Wallis test. Dunn's multiple comparison test with Benjamini-Hochberg correction was subsequently used for post-hoc analysis.

### Supplementary Fig. 1

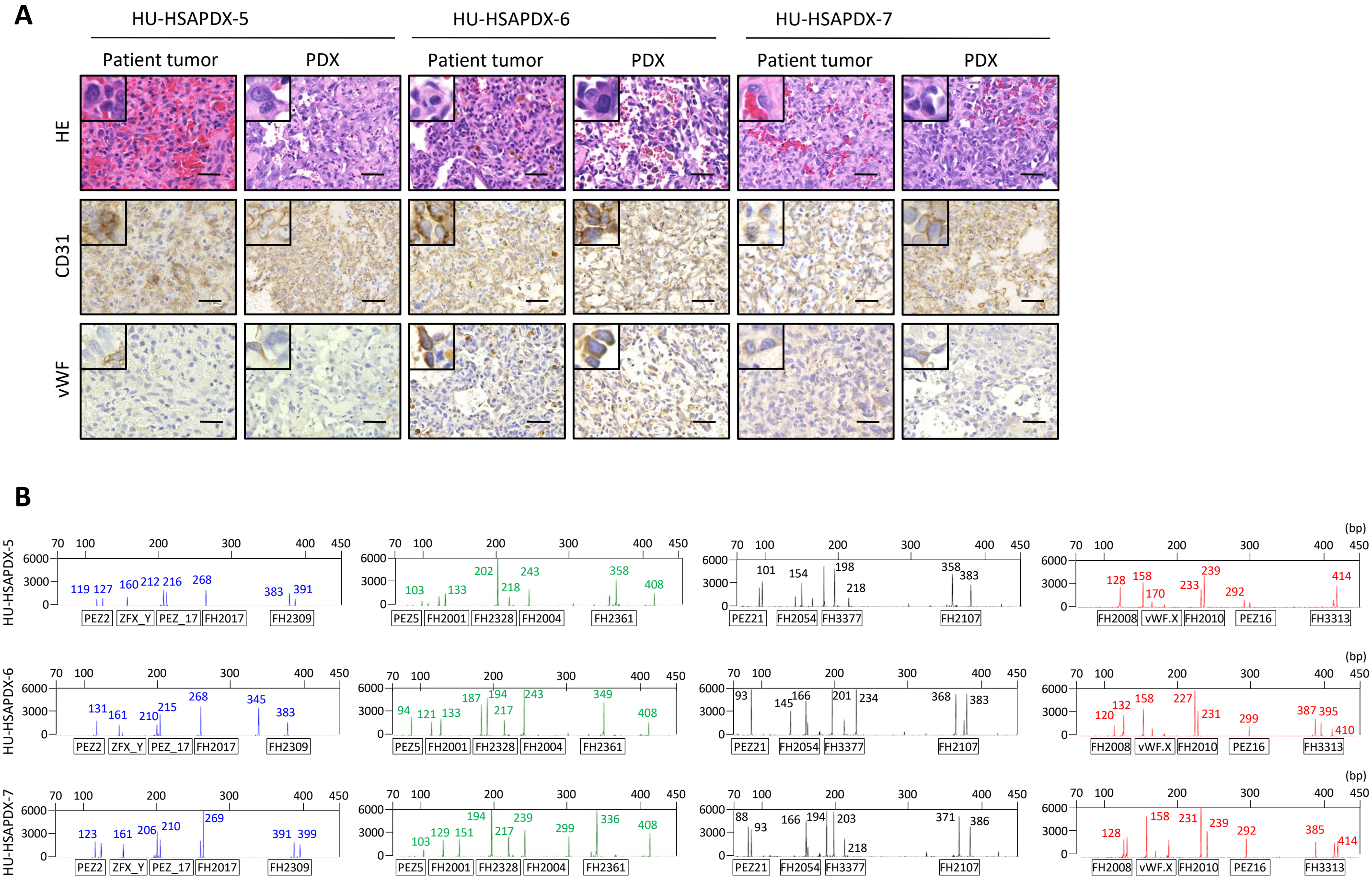


**Fig. S1. Establishment of canine HSA patient-derived xenograft (PDX) models.**
**(A)** Representative images of H&E staining and IHC for the endothelial markers CD31 and vWF. Images compare the original patient tumor with the corresponding PDX models. Insets show magnified views. Scale bars = 50 µm.
**(B)** STR profiles of the established PDX models.

| Protein Name | Manufacturer | Host | Usage | Dilution | Catalog no. |
|---|---|---|---|---|---|
| LDHA | Homemade | Rabbit | WB | 1:1,000 | - |
| LDHB | Santa Cruz Biotechnology | Mouse | WB | 1:1,000 | sc-100775 |
| SREBP2 | Santa Cruz Biotechnology | Mouse | WB | 1:1,000 | sc-13552 |
| panKla | PTM BIO | Rabbit | WB | 1:1,000 | PTM-1401RM |
| pH3S10 | Cell Signaling Technology | Rabbit | WB | 1:1,000 | 3377 |
| Cleaved-caspase 3 | Cell Signaling Technology | Rabbit | WB | 1:1,000 | 9661S |
| Actin | Sigma-Aldrich | Mouse | WB | 1:10,000 | MAB1501 |
| H3 | MAB Institute | Mouse | WB | 1:10,000 | MABI0001-20 |
| Goat anti-Mouse IgG (H+L) | Thermo Fisher Scientific | Goat | WB | 1:10,000 | G21040 |
| Goat anti-Rabbit IgG (H+L) | Thermo Fisher Scientific | Goat | WB | 1:10,000 | G21234 |
| CD31 | Abcam | Mouse | IHC | 1:200 | ab134168 |
| vWF | Agilent technologies | Rabbit | IHC | 1:200 | A0082 |

Table S1. Antibody list

| Species | Target | | Sequence | Gene ID |
|---|---|---|---|---|
| Canine | *SQLE* | F | ACTTGGTGGCGAATGTGTTG | ENSCAFG00000001056 |
| | | R | GCTCGGGGTTTTGTAATCCAAG | |
| | *SREBF1* | F | ATTGAGAAGCGCTACC | ENSCAFG00000018436 |
| | | R | ATGTAATCGATGGCCT | |
| | *SREBF2* | F | TGAGTTGAAGGACCTGGTCATG | ENSCAFG00000001013 |
| | | R | TGTAGTCAATGGCCTTCCTCAG | |
| | *LDLR* | F | TTGCAAGGACAAGTCTGACG | ENSCAFG00000017539 |
| | | R | AGCCAATCTCATCGCTCATG | |
| | *HMGCS1* | F | TGCTAATTGGGCCAAATGCG | ENSCAFG00000018572 |
| | | R | TGCACTGAGGTAGCACTGTATG | |
| | *HMGCR* | F | ACTGCTGATGAAGGATCCAAGG | ENSCAFG00000009517 |
| | | R | ACAAAAAGGCCGTGCATTCG | |
| | *SCD* | F | GGTGGTTTGCTAGACTTGTCTG | ENSCAFG00000009663 |
| | | R | TCAACAGGTGCCAGGTTTG | |
| | *CYP51A1* | F | ACAGTCTGTGGAGAAGATCTGC | ENSCAFG00000001945 |
| | | R | CAATGGTCTGAGGAGTTTTGGC | |
| | *RPL32* | F | TGGTTACAGGAGCAACAAGAAA | ENSCAFG00000004871 |
| | | R | GCACATCAGCAGCACTTCA | |
| | *RPL13A* | F | GCCGGAAGGTTGTAGTCGT | ENSCAFG00000029892 |
| | | R | GGAGGAAGGCCAGGTAATTC | |
| | *TBP* | F | ATAAGAGAGCCCCGAACCAC | ENSCAFG00000004119 |
| | | R | TCACATCACAGCTCCCCAC | |
| | *YWHAZ* | F | TTACTTGGCCGAAGTTGCTG | ENSCAFG00000000580 |
| | | R | ACAGAGAAGTTAAGGGCCAGAC | |
| | *HMBS* | F | AGTCGACCTGGTTGTTCACTC | ENSCAFG00000012342 |
| | | R | AACAGCATCATAGGGGTTCTCC | |
| | *B2M* | F | ACGGAAAGGAGATGAAAGCA | ENSCAFG00000013633 |
| | | R | CCTGCTCATTGGGAGTGAA | |
| | *ACTB* | F | CCAGCAAGGATGAAGATCAAG | ENSCAFG00000016020 |
| | | R | TCTGCTGGAAGGTGGACAG | |

Table S2. Primer sequences for RT-qPCR

| Experiments | Reference genes | Results |
|---|---|---|
| RNA-seq results validation | *B2M, TBP* | Fig. 1F |
| (R)-GNE-140 treatment in HU-HSA-3 | *RPL32, HMBS* | Fig. 1F |
| Sodium oxamate treatment | *RPL32, YWHAZ* | Fig. 2C |
| LDHA or LDHB knockout | *HMBS, B2M* | Fig. 3B |

Table S3. Selected reference genes for qPCR

| Name | Sequences (5’ to 3’) |
| --- | --- |
| sgLDHA (F) | CACCGTACCTTCATTAAGATACTGA |
| sgLDHA (R) | AAACTCAGTATCTTAATGAAGGTAC |
| sgLDHB (F) | CACCGTGTTAATTAAATACCTGGGT |
| sgLDHB (R) | AAACACCCAGGTATTTAATTAACAC |
| sgScr (F) | CACCGATATACGATACTCCGTTGCC |
| sgScr (R) | AAACGGCAACGGAGTATCGTATATC |

Table S4. sgRNA sequences

| Patient ID | Breed | Age | Sex | Location |
| --- | --- | --- | --- | --- |
| HU-HSAPDX-3 | Flat Coated Retriever | 8y | Spayed female | Spleen |
| HU-HSAPDX-5 | Miniature Dachshund | 10y2m | Male | Spleen |
| HU-HSAPDX-6 | Chihuahua | 13y9m | Castrated male | Spleen |
| HU-HSAPDX-7 | Mix | 10y9m | Male | Spleen |

Table S5. Patient information of HSA PDX models
